# Pathology-Specific Modulation of Corticostriatal Circuitry by Chronic Alcohol Consumption in Alzheimer’s Disease Mouse Models

**DOI:** 10.1101/2025.08.15.670607

**Authors:** Yufei Huang, Xueyi Xie, Ruifeng Chen, Zhenbo Huang, Himanshu Gangal, Xuehua Wang, Jun Wang

**Author notes:** To whom correspondence should be addressed: Jun Wang, M.D., Ph.D., 8447 John Sharp Pkwy, 2106 Medical Research and Education Building Bryan, TX 77807. The authors declare no competing financial interests.

## Abstract

Chronic alcohol use is a major modifiable risk factor for Alzheimer’s disease (AD), yet the mechanisms by which it modulates AD pathophysiology remain unclear. Here, we examined circuit-level and pathological changes in two distinct AD mouse models, humanized Aβ knock-in (hAPP-KI) (Aβ-driven) and PS19 (tau-driven), subjected to a chronic intermittent alcohol exposure paradigm. In hAPP-KI mice, alcohol increased Aβ accumulation and excitatory transmission in the medial prefrontal cortex (mPFC) while reducing corticostriatal transmission and striatal cholinergic output. These alterations were accompanied by enhanced recruitment of microglia around Aβ plaques. In contrast, alcohol-exposed PS19 mice displayed elevated mPFC-to-dorsomedial striatum (DMS) glutamatergic transmission and increased tau phosphorylation without significant changes in microglial activation or local mPFC excitatory drive. In wild-type mice, microglial depletion enhanced glutamatergic transmission onto cortical neurons, suggesting a homeostatic role for microglia in maintaining excitatory balance. Together, these findings reveal pathology-specific effects of alcohol on circuit dysfunction and propose microglia as an important modulator of alcohol-induced synaptic remodeling in the early stage of AD.

**Highlights:**

1. Chronic alcohol exposure modulates glutamatergic transmission and neuroimmune responses in pathology-specific manners across Aβ- and tau-driven Alzheimer’s disease mouse models.
2. In hAPP-KI mice, alcohol increases mPFC excitatory input, Aβ plaque burden, and microglial activation, while impairing corticostriatal transmission and striatal cholinergic signaling.
3. In PS19 mice, alcohol enhances mPFC-to-striatum excitatory transmission and tau phosphorylation without altering local microglial activation or mPFC excitability.
4. Microglial depletion in wild-type mice recapitulates alcohol-induced glutamatergic changes in hAPP-KI mice, revealing a homeostatic role for microglia in regulating cortical excitatory balance.

## INTRODUCTION

Alzheimer’s disease (AD) is the most common form of dementia, affecting over 6 million Americans and more than 10% of individuals over age 65 (ASSOCIATION, 2023). It is characterized by progressive cognitive decline associated with two hallmark pathologies: the accumulation of extracellular β-amyloid (Aβ) and the intracellular hyperphosphorylation and spread of tau protein (Palop et al., 2007; Ittner and Gotz, 2011; Serrano-Pozo et al., 2011; Barnes et al., 2018; DeTure and Dickson, 2019; Breijyeh and Karaman, 2020). These pathologies emerge and progress in distinct brain regions: Aβ aggregation typically initiates in neocortical areas such as the mPFC, whereas tau pathology begins in the entorhinal cortex and hippocampus and later spreads to the neocortex (Palmqvist et al., 2017; Adams et al., 2019).

Beyond their molecular signatures, Aβ and tau exert distinct yet complementary effects on neural circuits. Aβ is associated with neuronal hyperexcitability and network dysfunction (Palop et al., 2007; Busche Marc et al., 2008; Zott et al., 2018; Zott et al., 2019; Guo et al., 2022). Among the affected pathways, the corticostriatal projection from the mPFC to the DMS is critical for goal-directed behavior, cognitive flexibility, and behavioral inhibition—functions that are impaired early in AD (Huang et al., 2025). In contrast, tau pathology is linked to synaptic degeneration, circuit hypoactivity, and progressive disconnection across brain networks, features that are more prominent in later-stage disease (Busche et al., 2019; Busche and Hyman, 2020; Harris et al., 2025).

Chronic alcohol consumption is a modifiable risk factor for cognitive decline and dementia and is known to disrupt cholinergic signaling (Lees et al., 2020; Ma et al., 2022; Huang et al., 2024). Alcohol also alters glutamatergic transmission, induces synaptic remodeling, and impairs neural circuits involved in executive function and behavioral regulation, particularly within corticostriatal pathways (Ma et al., 2017; Ma et al., 2018; Lu et al., 2019; Gangal et al., 2023; Xie et al., 2023). While the molecular hallmarks of AD have been extensively studied, there is increasing recognition that environmental and lifestyle factors such as alcohol use can accelerate disease progression and increase the risk of cognitive impairment (Rehm, 2011; Heymann et al., 2016; Sabia et al., 2018; Tousley et al., 2023; Fisher et al., 2025; Kang et al., 2025). However, the circuit-level mechanisms through which alcohol interacts with Aβ and tau pathology remain poorly understood.

To address this gap, we examined how chronic intermittent alcohol exposure influences corticostriatal glutamatergic transmission in multiple AD mouse models. Our findings reveal that alcohol differentially alters corticostriatal synaptic transmission and microglial responses depending on the underlying pathology. By directly comparing Aβ- and tau-driven models, we identify pathology-specific mechanisms through which alcohol impacts neural circuit function, providing new insight into how environmental factors can exacerbate AD progression and highlighting potential circuit-level targets for therapeutic intervention.

## RESULTS

### hAPP KI mice exhibited hyperactive locomotor activity during withdrawal from chronic alcohol consumption

To examine how chronic alcohol intake influences AD-related pathology, we used an intermittent two-bottle choice (2BC) drinking paradigm. Beginning at 2 months of age, mice were given access to 20% alcohol every other day for 16 weeks. Twenty-four hours after the final drinking session, locomotor activity was assessed to determine the behavioral effects of prolonged alcohol exposure and withdrawal (Fig. 1A). During the 2BC procedure, five sessions of single housing were used to quantify individual alcohol and water consumption. Water intake did not differ significantly between genotypes (Fig. 1B, *F*_(1,13)_ = 2.952, *p* = 0.1095), whereas hAPP-KI mice showed a marginally higher alcohol intake than wild-type (WT) mice (Fig. 1C, *F*_(1,13)_ = 3.79, *p* = 0.0734). However, alcohol preference was similar between the two groups (Fig. 1D, *F*_(1,13)_ = 0.009, *p* = 0.924), suggesting that the slight increase in alcohol intake in hAPP-KI mice was not due to altered reward sensitivity. To explore whether activity levels contributed to these differences, we measured spontaneous locomotor activity in both water- and alcohol-drinking mice. Under water-drinking conditions, hAPP-KI and WT mice exhibited comparable locomotor activity (Fig. 1E, *F*_(1,13)_ = 0.096, *p* = 0.761; Fig. 1F, t_13_ = 0.3102, *p* = 0.7613). In contrast, following chronic alcohol exposure, hAPP-KI mice displayed significantly greater locomotor activity than WT mice 24 hours after the final drinking session (Fig. 1G, *F*_(1,13)_ = 17.94, *p* = 0.001 and Fig. 1H, t_13_ = 4.235, *p* = 0.001). These findings indicate that chronic intermittent alcohol exposure, followed by withdrawal, selectively increases locomotor activity in hAPP-KI mice compared with WT controls.

**Figure 1.**
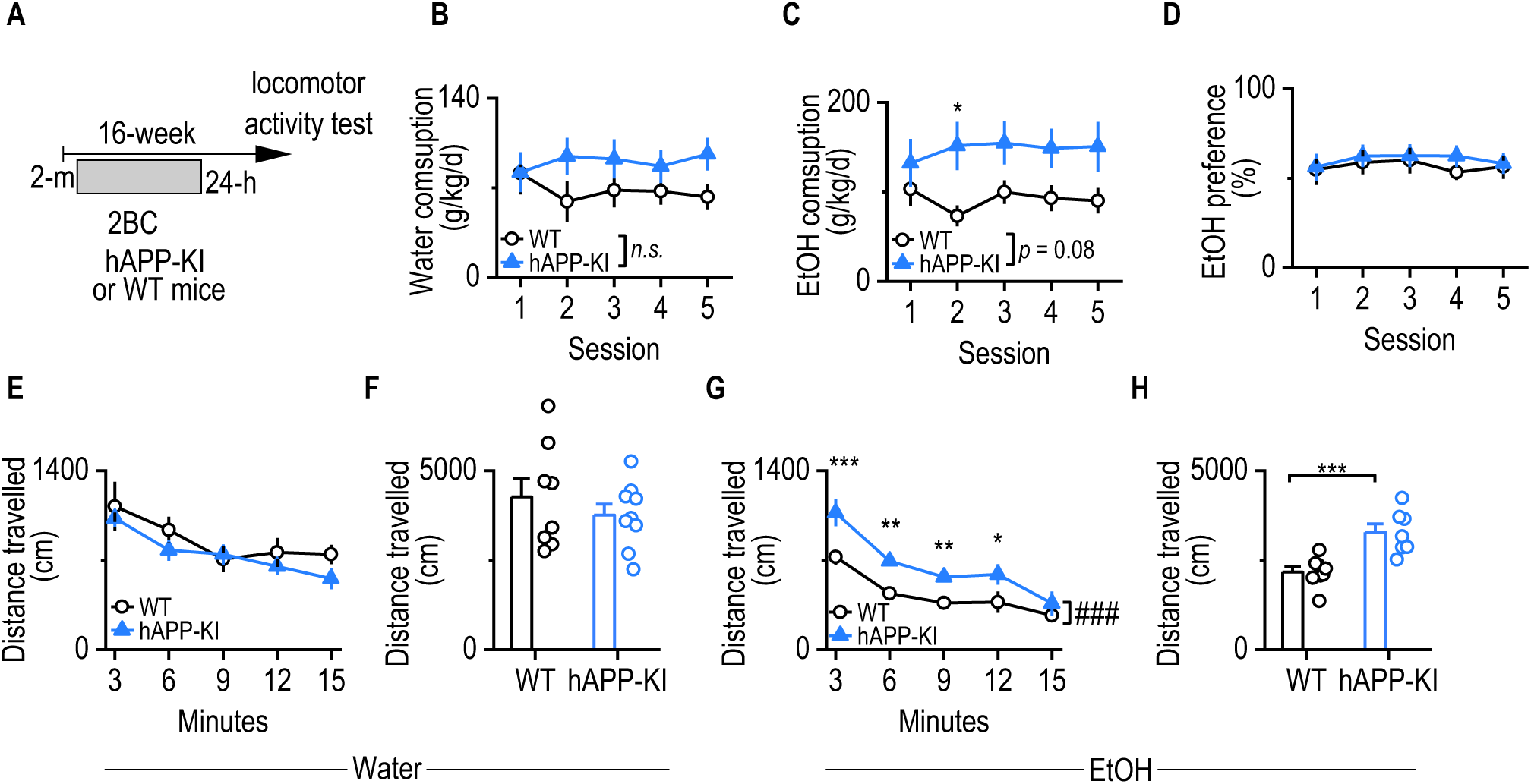
hAPP KI mice exhibited hyperactive locomotor activity during withdrawal from chronic alcohol consumption. **A**, Schematic of the experimental timeline. Two-month-old wild-type (WT) and hAPP knock-in (hAPP-KI) mice were subjected to 16 weeks of intermittent access two-bottle choice (2BC) drinking with 20% alcohol (EtOH), followed by a 24-hour withdrawal period before assessment of locomotor activity in the open field test. **B**, Water intake (g/kg/day) across five drinking sessions was comparable between WT and hAPP-KI mice (n.s.). Two-way repeated-measures (RM) ANOVA with Greenhouse–Geisser correction followed by Sidak’s post hoc test. n = 8 (WT) and 7 (hAPP-KI). **C**, Alcohol intake (g/kg/day) did not significantly differ between groups across sessions. Two-way RM ANOVA with Greenhouse–Geisser correction followed by Sidak’s post hoc test. n = 8 (WT) and 7 (hAPP-KI) mice. **D,** Alcohol preference was similar between WT and hAPP-KI mice. Two-way RM ANOVA with Greenhouse–Geisser correction followed by Sidak’s post hoc test. n = 8 (WT) and 7 (hAPP-KI) mice. **E, F,** Locomotor activity following water intake did not differ between WT and hAPP-KI mice (E: time course; F: total distance traveled). Two-way RM ANOVA with Greenhouse–Geisser correction and Sidak’s post hoc test (E); unpaired t-test (F). n = 6 (WT) and 9 (hAPP-KI) mice. **G**, **H,** Alcohol-exposed hAPP-KI mice showed significantly elevated locomotor activity during withdrawal compared to WT controls (G: time course; H: total distance traveled). Two-way RM ANOVA with Greenhouse–Geisser correction and Sidak’s post hoc test (G); unpaired t-test (H). n = 8 (WT) and 7 (hAPP-KI) mice. \**p* < 0.05, \*\**p* < 0.01.

### Chronic alcohol consumption increased glutamatergic inputs and Aβ accumulation in the mPFC in hAPP-KI mice

Previous studies have shown that mPFC neurons exhibit hyperactivity and enhanced glutamatergic input during the early stages of AD(Huang et al., 2025). To determine whether chronic alcohol consumption exacerbates these neurophysiological changes and alters Aβ pathology, hAPP-KI mice were subjected to a 16-week intermittent 2BC drinking paradigm. Electrophysiological recordings were performed on mPFC slices 24 hours after the final alcohol exposure (Fig. 2A).

**Figure 2.**
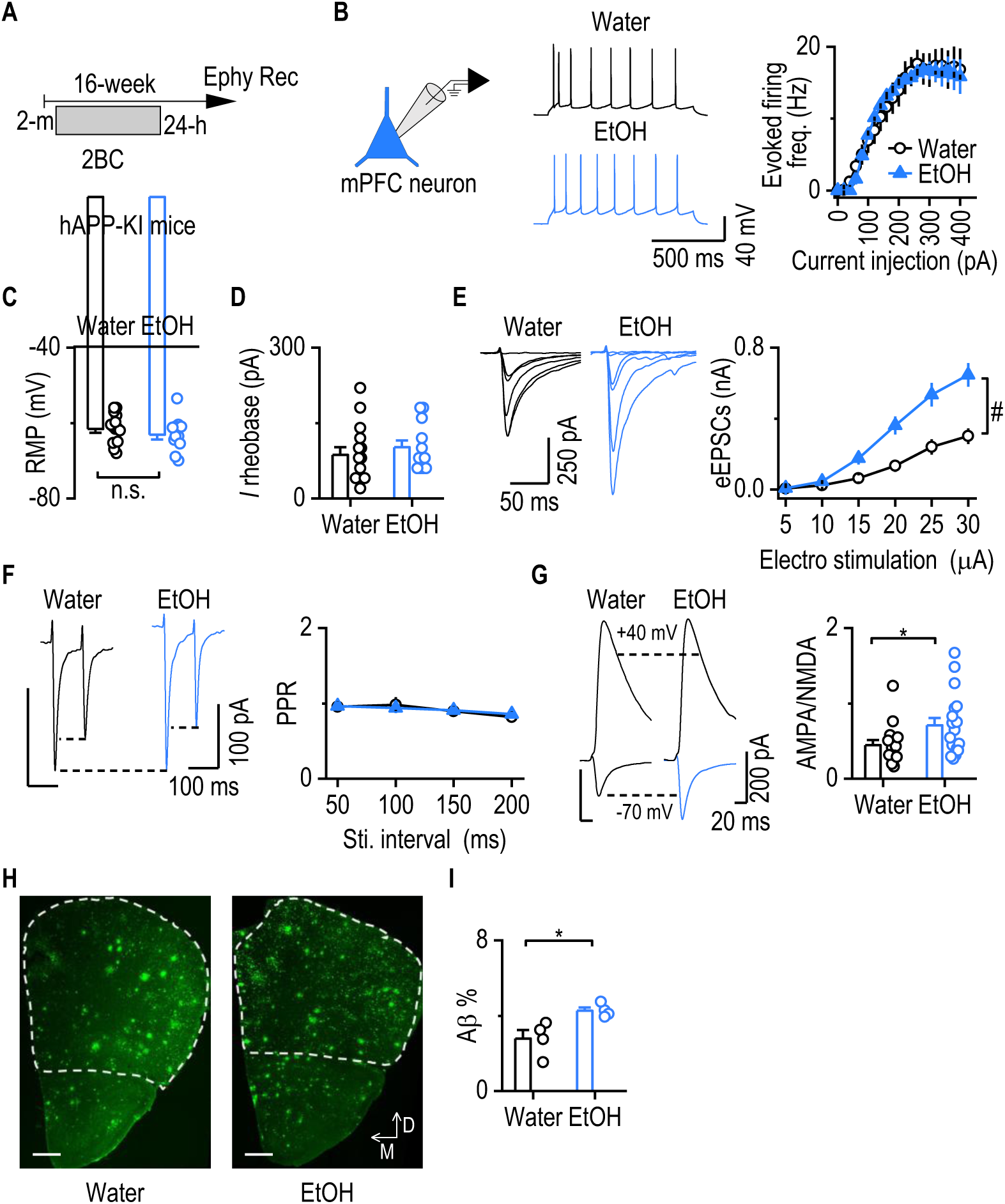
Chronic alcohol consumption increases glutamatergic transmission and Aβ accumulation in the mPFC of hAPP-KI mice. **A,** Schematic of the 2BC paradigm in hAPP-KI mice. Mice underwent 16 weeks of alcohol exposure starting at 2 months of age. Electrophysiological recordings were performed 24 hours after the final alcohol session. **B,** Diagram showing patch-clamp recording in mPFC neurons and sample traces of evoked firing. Neuronal excitability in the mPFC neurons did not differ between water- and alcohol-drinking hAPP-KI mice. Two-way RM ANOVA with Greenhouse–Geisser correction and Sidak’s post hoc test. n = 13 neurons from 3 mice (13/3) for the Water group and 8/3 for the EtOH group. **C,** Resting membrane potential (RMP) of mPFC neurons was similar between groups. Unpaired t-test. n = 15/3 (Water) and 12/3 (EtOH). **D,** Rheobase current did not differ between water- and alcohol-drinking hAPP-KI mice. Unpaired t-test. n = 15/3 (Water) and 12/3 (EtOH). **E,** Amplitudes of electrically evoked excitatory postsynaptic currents (eEPSCs) were significantly higher in alcohol-drinking hAPP-KI mice compared to water-drinking controls. Two-way RM ANOVA with Greenhouse–Geisser correction and Sidak’s post hoc test. n = 15/3 (Water) and 20/4 (EtOH). **F,** Paired-pulse ratio (PPR) of eEPSCs did not differ between groups. Two-way RM ANOVA with Greenhouse–Geisser correction and Sidak’s post hoc test. n = 13/3 (Water) and 20/4 (EtOH). **G,** AMPA/NMDA ratio was significantly higher in alcohol-drinking hAPP-KI mice than in water-drinking mice. Mann Whitney test. n = 15/3 (Water) and 18/4 (EtOH). **H,** Representative images of Aβ staining in the cortical region. Scale bar: 500 μm. **I,** Aβ-covered cortical area was significantly larger in alcohol-drinking hAPP-KI mice. Unpaired t-test. n = 4 mice from each group.

Compared with water-drinking hAPP-KI mice, alcohol-exposed mice showed no significant differences in evoked firing rates (Fig. 2B, *F*_(1,19)_ = 0.028, *p* = 0.8689), resting membrane potential (Fig. 2C, t_25_ = 0.9672, *p* = 0.3427), or rheobase current (Fig. 2D, t_25_ = 0.7252, *p* = 0.4751), indicating that overall neuronal excitability was not altered by alcohol consumption. In contrast, alcohol-exposed hAPP-KI mice exhibited significantly increased electrically evoked excitatory postsynaptic currents (eEPSCs) in mPFC neurons (Fig. 2E, *F*_(1,33)_ = 13.52, *p* = 0.0008), suggesting enhanced excitatory synaptic input. Paired-pulse ratios (PPRs) were unchanged (Fig. 2F, *F*_(1,31)_ = 0.002, *p* = 0.9649), indicating that presynaptic glutamate release probability was unaffected. Consistent with the increased eEPSCs, the AMPA/NMDA ratio was significantly higher in the alcohol group compared with the water group (Fig. 2G, *U* = 76, *p* = 0.0331), further supporting a postsynaptic enhancement of AMPA-induced excitatory transmission.

To assess whether alcohol affected Aβ pathology, we performed immunostaining and quantified the Aβ plaque-covered area in the cortex (Fig. 2H). Alcohol-drinking hAPP-KI mice showed a significantly greater Aβ-covered area than water-drinking controls (Fig. 2I, t_6_ = 3.029, *p* = 0.0231), indicating that chronic alcohol intake promotes Aβ accumulation in the cortical region.

Together, these findings suggest that chronic alcohol consumption enhances excitatory synaptic transmission in the mPFC and accelerates Aβ pathology in hAPP-KI mice.

### Chronic alcohol consumption decreased glutamatergic mPFC to DMS transmission and striatal ACh levels in hAPP-KI mice

Previous studies have reported that alcohol exposure potentiates excitatory inputs from the mPFC to the DMS (Ma et al., 2017; Ma et al., 2018; Lu et al., 2019; Xie et al., 2023), and that corticostriatal circuits exhibit hyperactivity during the early stages of AD (Huang et al., 2025). To investigate how chronic alcohol consumption affects corticostriatal transmission in an Aβ pathology model, we infused AAV-Chronos-GFP into the mPFC of hAPP-KI mice (Fig. 3A) and subjected them to a 4-month intermittent 2BC drinking paradigm. Following the final drinking session, mPFC-to-DMS synaptic transmission was assessed by optogenetically stimulating mPFC terminals with 470 nm light while recording from medium spiny neurons (MSNs) in the DMS (Fig. 3B). Compared with water-drinking controls, alcohol-exposed hAPP-KI mice exhibited significantly reduced optogenetically evoked EPSC (oEPSC) amplitudes in MSNs (Fig. 3B, *F*_(1,36)_ = 5.204, *p* = 0.0286), indicating diminished corticostriatal excitatory input. The PPRs were significantly increased in the alcohol group(Fig. 3C, *F*_(1,31)_ = 4.336, *p* = 0.0457), suggesting reduced presynaptic glutamate release probability. In addition, the AMPA/NMDA ratio was significantly lower in alcohol-exposed mice than in water-drinking controls (Fig. 3D, *U* = 72, *p* = 0.0207), consistent with a reduction in AMPA receptor–mediated transmission. Together, these findings indicate that chronic alcohol consumption attenuates mPFC-to-DMS excitatory synaptic input in hAPP-KI mice.

**Figure 3.**
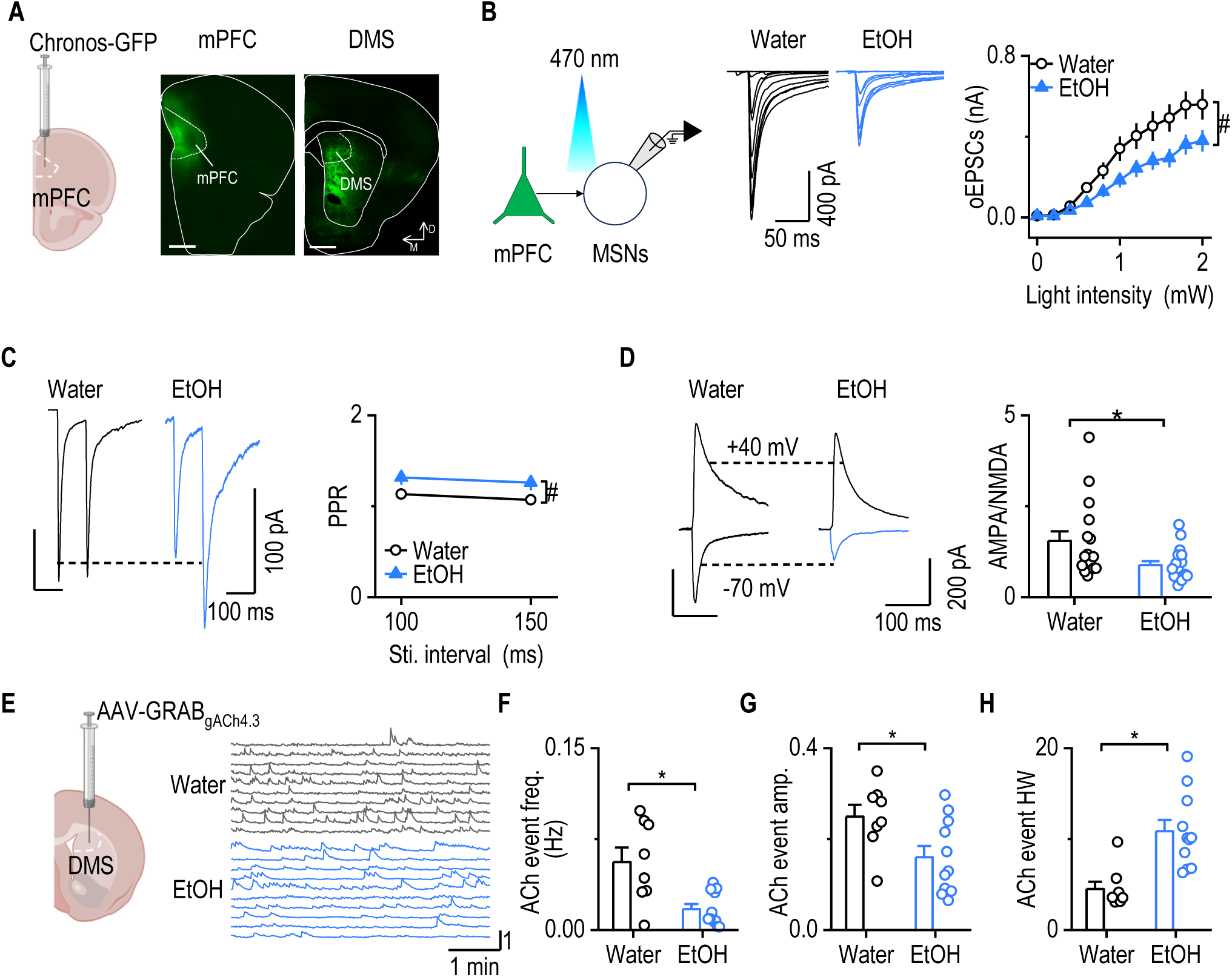
Chronic alcohol consumption reduces mPFC-to-DMS glutamatergic transmission and striatal ACh levels in hAPP-KI mice. **A,** Schematic illustrating AAV-Chronos-GFP injection into the medial prefrontal cortex (mPFC) and expression in both the mPFC and dorsomedial striatum (DMS). Scale bar: 500 μm. **B**, Optogenetically evoked glutamatergic transmission from mPFC to DMS medium spiny neurons (MSNs) was significantly reduced in alcohol-drinking hAPP-KI mice compared to water controls. Two-way repeated-measures (RM) ANOVA with Greenhouse–Geisser correction, followed by Sidak’s post hoc test. n = 18 neurons from 3 mice (18/3, Water), 20 neurons from 4 mice (20/4, EtOH). **C**, Paired-pulse ratios (PPRs) of optogenetically evoked EPSCs were significantly higher in alcohol-drinking mice, suggesting reduced presynaptic release probability. Two-way RM ANOVA with Greenhouse–Geisser correction, followed by Sidak’s post hoc test; ^#^p < 0.05. n = 16/3 (Water), 17/4 (EtOH). **D,** The AMPA/NMDA ratio was significantly lower in alcohol-drinking mice than in water controls. Mann Whitney test, \**p* < 0.05. n = 16 neurons from 3 mice (16/3, Water), 17 neurons from 4 mice (17/4, EtOH). **E**, Representative traces of spontaneous striatal acetylcholine (ACh) release events. **F**, Frequency of spontaneous ACh release events was significantly lower in alcohol-drinking hAPP-KI mice. Mann Whitney test, \**p* < 0.05. n = 8 slices from 3 mice (8/3, Water), 11 slices from 4 mice (11/4, EtOH). **G,** The amplitude of spontaneous ACh release events was significantly reduced in alcohol-drinking mice. Unpaired t-test, \**p* < 0.05. n = 8 slices from 3 mice (8/3, Water), 11 slices from 4 mice (11/4, EtOH). **H**, Half-width (HW) of spontaneous ACh events was significantly increased in the alcohol group. Mann Whitney test, \*\*\**p* < 0.001. n = 8 slices from 3 mice (8/3, Water), 11 slices from 4 mice (11/4, EtOH).

Given prior evidence that alcohol consumption disrupts cholinergic signaling (Ma et al., 2022; Huang et al., 2024), we next examined striatal acetylcholine (ACh) dynamics. A genetically encoded ACh sensor was expressed in the dorsomedial striatum (Fig. 3E). In alcohol-exposed hAPP-KI mice, the frequency (Fig. 3F, *U* = 15, *p* = 0.0157) and amplitude of spontaneous ACh release events (Fig. 3G, t_17_ = 2.458, *p* = 0.025) were significantly reduced. Notably, the half-width of ACh transients was increased (Fig. 3H, *U* = 4, *p* = 0.0003), indicating impaired ACh clearance and altered cholinergic tone.

In summary, chronic alcohol consumption reduces excitatory input from the mPFC to the DMS and impairs striatal cholinergic dynamics in hAPP-KI mice.

### Chronic alcohol consumption increased glutamatergic mPFC to DMS transmission in PS19 mice

While Aβ accumulation characterizes the early stages of AD, pathological tau hyperphosphorylation and spreading define later stages and are strongly associated with widespread neuronal dysfunction and cognitive decline (Busche and Hyman, 2020; Knopman et al., 2021; Harris et al., 2025). To investigate how alcohol consumption influences neural circuit function under tau pathology, we first examined the impact of tau overexpression on corticostriatal transmission.

We co-injected AAV-ChrimsonR-tdTomato and either AAV-Tau-GFP or control AAV-GFP into the mPFC to enable optogenetic activation of mPFC terminals in the striatum while inducing tau expression in mPFC neurons (Fig. 4A). Histological analysis confirmed robust phospho-tau expression in the mPFC following AAV-Tau-GFP infusion, whereas AAV-GFP controls showed no detectable phospho-tau (Fig. 4B). Optogenetic stimulation of mPFC terminals in the DMS revealed that tau-expressing mice exhibited significantly reduced oEPSC amplitudes (Fig. 4C and D, *F*_(1,30)_ = 8.209, *p* = 0.0075) and increased PPRs (Fig. 4E, *F*_(1,19)_ = 14.39, *p* = 0.0012), suggesting impaired corticostriatal transmission likely due to presynaptic dysfunction.

**Figure 4.**
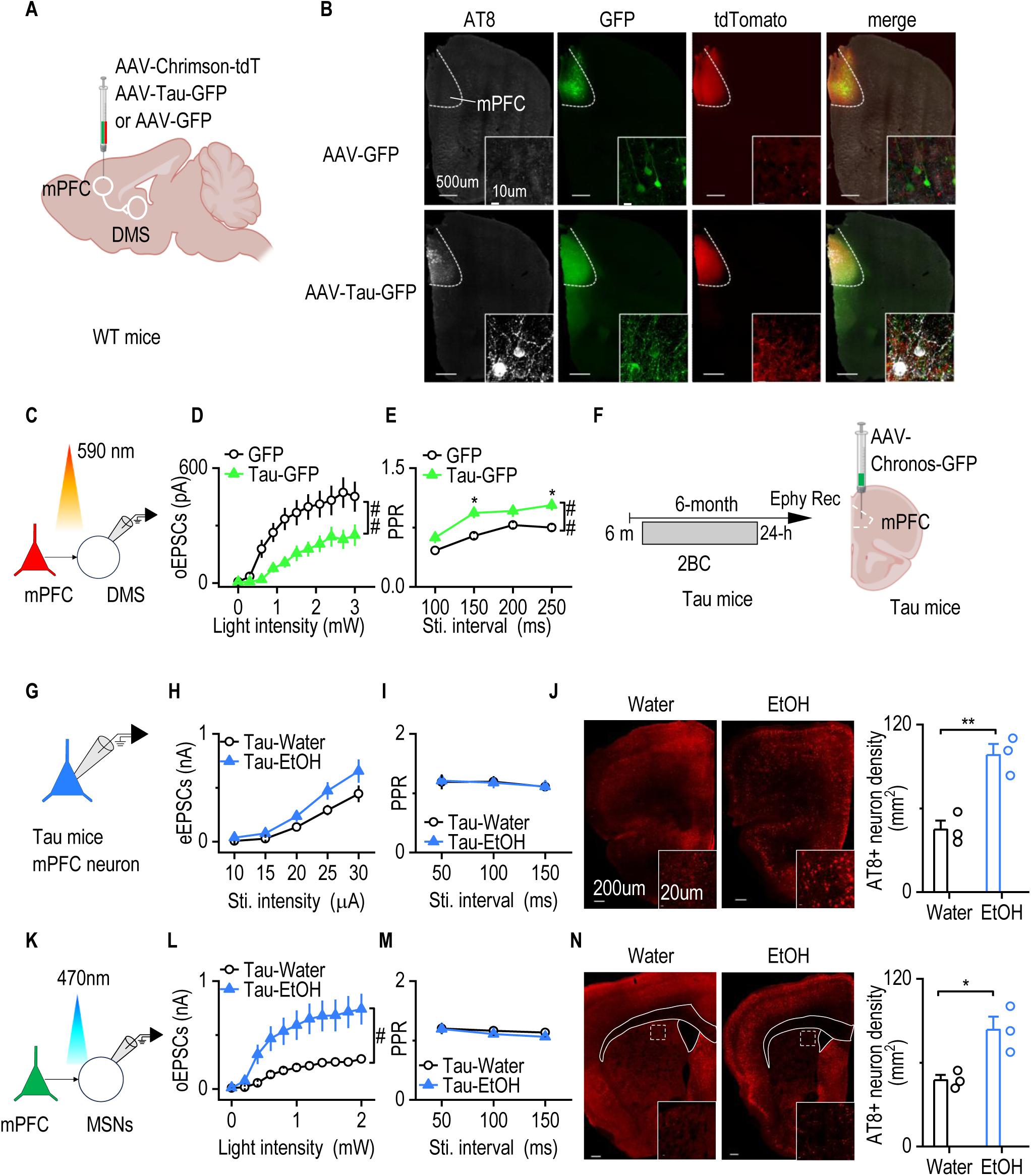
Chronic alcohol consumption enhances mPFC-to-DMS glutamatergic transmission in PS19 mice, a tauopathy model. **A,** Schematic of the viral injection strategy in wild-type (WT) mice. AAV-Chrimson and AAV-Tau-GFP or AAV-GFP (control) were co-infused into the medial prefrontal cortex (mPFC). **B**, Representative images showing AT8 immunostaining and co-expression of GFP and Chrimson in the mPFC. Scale bar: 500 μm; 10 μm for the enlarged image. **C**, Schematic of oEPSC recordings from DMS MSNs following mPFC terminal stimulation. **D**, oEPSC amplitudes were significantly lower in AAV-Tau-GFP–injected mice compared to GFP controls. Two-way RM ANOVA with Greenhouse–Geisser correction, followed by Sidak’s post hoc test; ^#^p < 0.05. n = 16 neurons from 3 (16/3) GFP- or Tau-injected mice. **E,** PPRs of oEPSCs were significantly higher in AAV-Tau-GFP–injected mice. Two-way RM ANOVA with Greenhouse–Geisser correction, followed by Sidak’s post hoc test; ^#^p < 0.05. n = 11/3 (GFP), 10/4 (Tau). **F,** Schematic of the 2BC alcohol exposure paradigm in PS19 mice. Mice underwent 6 months of alcohol consumption starting at 6 months of age. AAV-Chronos-GFP was infused into the mPFC for mPFC-to-DMS transmission recording. Electrophysiological recordings were performed 24 hours after the final session. **G,** Diagram illustrating whole-cell patch-clamp recordings of mPFC neurons in PS19 mice. **H,** Electrically evoked EPSC (eEPSC) amplitudes in mPFC neurons were not different between water- and alcohol-drinking PS19 mice. Two-way RM ANOVA with Greenhouse–Geisser correction, followed by Sidak’s post hoc test; ^#^p < 0.05. n = 12/3 (Water), 13/4 (EtOH). **I**, PPRs of eEPSCs in mPFC neurons were similar between the two groups. Two-way RM ANOVA with Greenhouse–Geisser correction, followed by Sidak’s post hoc test; ^#^p < 0.05. n = 16/3 (Water), 17/4 (EtOH). **J,** Sample images of AT8+ neurons in the mPFC of water and EtOH groups. The density of AT8+ neurons is significantly higher in the EtOH group than in the water group. Unpaired t-test, \*\**p* < 0.01. n = 3 mice from each group. **K,** Diagram showing optogenetic stimulation of mPFC terminals and patch-clamp recordings from DMS MSNs. **L,** oEPSC amplitudes in DMS MSNs were significantly increased in alcohol-drinking PS19 mice compared to water controls. Two-way RM ANOVA with Greenhouse–Geisser correction, followed by Sidak’s post hoc test; ^#^p < 0.05. n = 10/3 (Water), 11/4 (EtOH). **M**, PPRs of oEPSCs were not significantly different between alcohol and water groups in PS19 mice. Two-way RM ANOVA with Greenhouse–Geisser correction, followed by Sidak’s post hoc test; ^#^p < 0.05. n = 12/3 (Water), 13/4 (EtOH). **N,** Sample images of AT8+ neurons in the DMS of water and EtOH groups. The density of AT8+ neurons is significantly higher in the EtOH group than in the water group. Unpaired t-test, \**p* < 0.05. n = 3 mice from each group.

To further examine the combined effects of Aβ and tau pathologies, we crossed PS19 tau mice with hAPP-KI mice to generate Tau;hAPP-KI double-transgenic mice (Sup. Fig. 1A). In these mice, the amplitude of eEPSCs in mPFC neurons was significantly lower than in hAPP-KI mice (Sup. Fig. 1B, F_(1,18)_ = 5.782, *p* = 0.0272) and PPRs were increased (Sup. Fig. 1C, F_(1,23)_ = 6.922, *p* = 0.0149), indicating reduced presynaptic glutamatergic transmission. Similarly, eEPSC amplitudes recorded from DMS MSNs were significantly reduced in Tau;hAPP-KI mice compared with hAPP-KI controls (Sup. Fig. 1D, F_(1,26)_ = 4.883, *p* = 0.0361), although PPRs were unchanged (Sup. Fig. 1E, F_(1,27)_ = 0.4894, *p* = 0.4902). In addition, MSNs from Tau;hAPP-KI mice exhibited reduced excitability and increased rheobase current (Sup. Fig. 1F, F_(1,19)_ = 14.82, *p* = 0.0011 and G, t_19_ = 2.921, *p* = 0.088), suggesting that tau expression dampens both synaptic input and neuronal output. Collectively, these results indicate that tau pathology disrupts cortical and striatal synaptic transmission as well as striatal excitability.

To evaluate how chronic alcohol exposure interacts with tau pathology, we subjected PS19 mice expressing the P301S mutant human tau transgene to a 6-month intermittent 2BC alcohol-drinking paradigm (Fig. 4F). Electrophysiological recordings were conducted 24 hours after the final drinking session. Alcohol-drinking PS19 mice showed no significant differences in eEPSC amplitude (Fig. 4G and H, *F*_(1,23)_ = 3.459, *p* = 0.0757) or PPRs (Fig. 4I, *F*_(1,22)_ = 0.0005, *p* = 0.9826) in mPFC neurons compared with water-drinking PS19 mice, indicating that local excitatory input to mPFC neurons was unaffected by alcohol under tau pathology. However, alcohol consumption significantly increased phosphorylated tau levels in the mPFC, as evidenced by higher densities of AT8-positive neurons in alcohol-exposed mice (Fig. 4J, t_4_ = 5.528, *p* = 0.0063).

To assess long-range corticostriatal connectivity, we recorded oEPSCs in the DMS following mPFC terminal stimulation (Fig. 4K). Alcohol-exposed PS19 mice displayed significantly higher oEPSC amplitudes compared with water-drinking controls (Fig. 4L, *F*_(1,19)_ = 8.036, *p* = 0.0106), while PPRs remained unchanged (Fig. 4M, *F*_(1,23)_ = 0.2573, *p* = 0.6168). The density of AT8-positive neurons in the DMS was also significantly increased in the alcohol group (Fig. 4N, t_4_ = 3.543, *p* = 0.024), suggesting a system-wide enhancement of tau phosphorylation following alcohol exposure.

Given that PS19 mice model tauopathy in isolation, we next asked how alcohol consumption influences corticostriatal transmission in the presence of both Aβ and tau pathologies, which could reveal whether one pathology predominates in shaping these changes. To address this, we crossed 5xFAD mice with PS19 mice to generate the 5xFAD;PS19 line, which exhibits both Aβ and tau pathologies. We next examined whether alcohol modifies synaptic impairments in 5xFAD;Tau mice. In these double-transgenic mice, eEPSC amplitudes recorded from mPFC neurons were comparable to those observed in PS19 mice following alcohol exposure (Sup. Fig. 2A, F_(1,20)_ = 0.1124, *p* = 0.7409), suggesting that local excitatory input within the mPFC was preserved. In contrast, optogenetically evoked mPFC-to-DMS transmission was significantly reduced in alcohol-exposed 5xFAD;Tau mice compared with PS19 controls (Sup. Fig. 2B, F_(1,16)_ = 9.494, *p* = 0.0072), indicating the presence of Aβ pathology limits the synaptic facilitation effect of alcohol seen in tau-only models, particularly at projection-specific synapses.

In summary, chronic alcohol consumption enhances mPFC-to-DMS glutamatergic transmission and increases tau phosphorylation in PS19 mice, whereas the presence of Aβ pathology blunts these alcohol-induced synaptic effects and further disrupts corticostriatal signaling.

### Chronic alcohol consumption increased microglia activation in mPFC in hAPP-KI mice

Microglia are the primary immune cells of the central nervous system and play essential roles in regulating neuronal activity (Badimon et al., 2020; Lv et al., 2024). To examine how alcohol consumption influences microglial responses in AD-related pathologies, we performed co-immunostaining for microglia and Aβ or tau in hAPP-KI and PS19 mice.

In hAPP-KI mice, alcohol-drinking animals exhibited a significantly higher density of microglia surrounding Aβ plaques in the mPFC compared with water-drinking controls (Fig. 5A and B, t_6_ = 3.154, *p* = 0.0197), indicating enhanced microglial recruitment in response to alcohol in the context of Aβ pathology.

**Figure 5.**
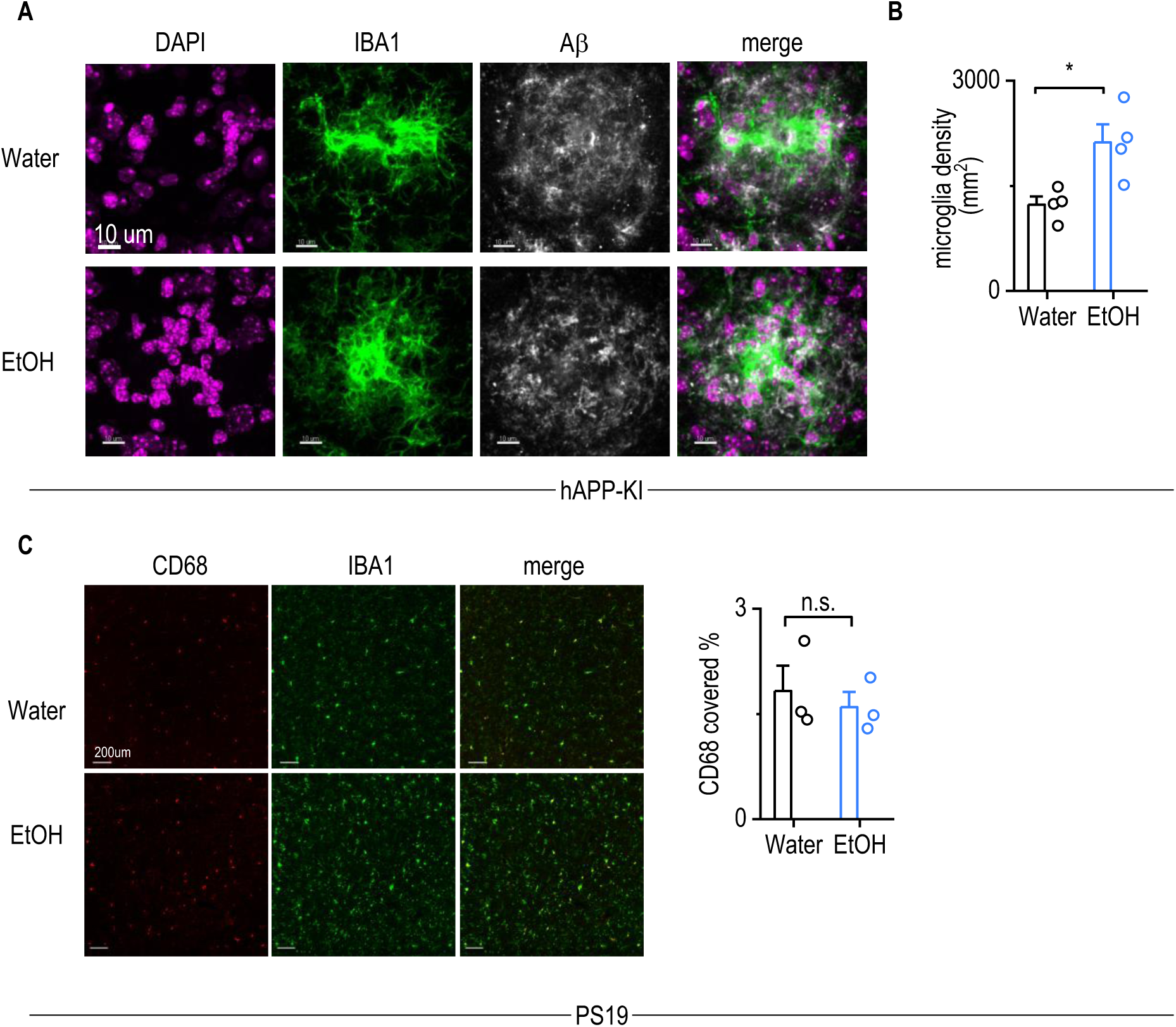
Chronic alcohol consumption increases microglial activation in the mPFC of hAPP-KI mice. **A,** Representative images showing microglia surrounding Aβ plaques in the cortical region of water- and alcohol-drinking hAPP-KI mice. **B,** The number of activated microglia surrounding Aβ plaques was significantly higher in alcohol-drinking hAPP-KI mice than water-drinking controls. Unpaired t-test, \**p* < 0.05. n = 4 mice from each group. **C**, Representative images of IBA1 and CD68 staining in the mPFC of water- and alcohol-drinking PS19 mice. No significant difference between water and EtOH groups in CD68-covered area. Unpaired t-test. n = 3 mice from each group.

To assess microglial activation in the PS19 tauopathy model, we stained for CD68, a marker of activated microglia (Fig. 5C). In contrast to the findings in hAPP-KI mice, alcohol consumption did not significantly alter CD68 expression in the mPFC of PS19 mice (Fig. 5C, t_4_ = 0.5547, *p* = 0.6086), suggesting that alcohol has minimal effects on microglial activation in the presence of tau pathology.

### Glutamatergic transmission is enhanced in microglia-deficient conditions in WT mice

Activated microglia enhance phagocytic capacity and promote synapse elimination; however, excessive activation can trigger cytokine release and neurotoxicity, ultimately contributing to neuronal apoptosis (Lv et al., 2024). In contrast, resting microglia help preserve network stability by regulating synchronized neuronal firing and modulating synaptic function(Badimon et al., 2020). To further investigate the role of microglia in regulating excitatory synaptic transmission, we depleted microglia in WT mice using the CSF1R inhibitor PLX5622 administered via intraperitoneal injection for 7 days, followed by electrophysiological recordings (Fig. 6A) as previously described (Gangal et al., 2025).

**Figure 6.**
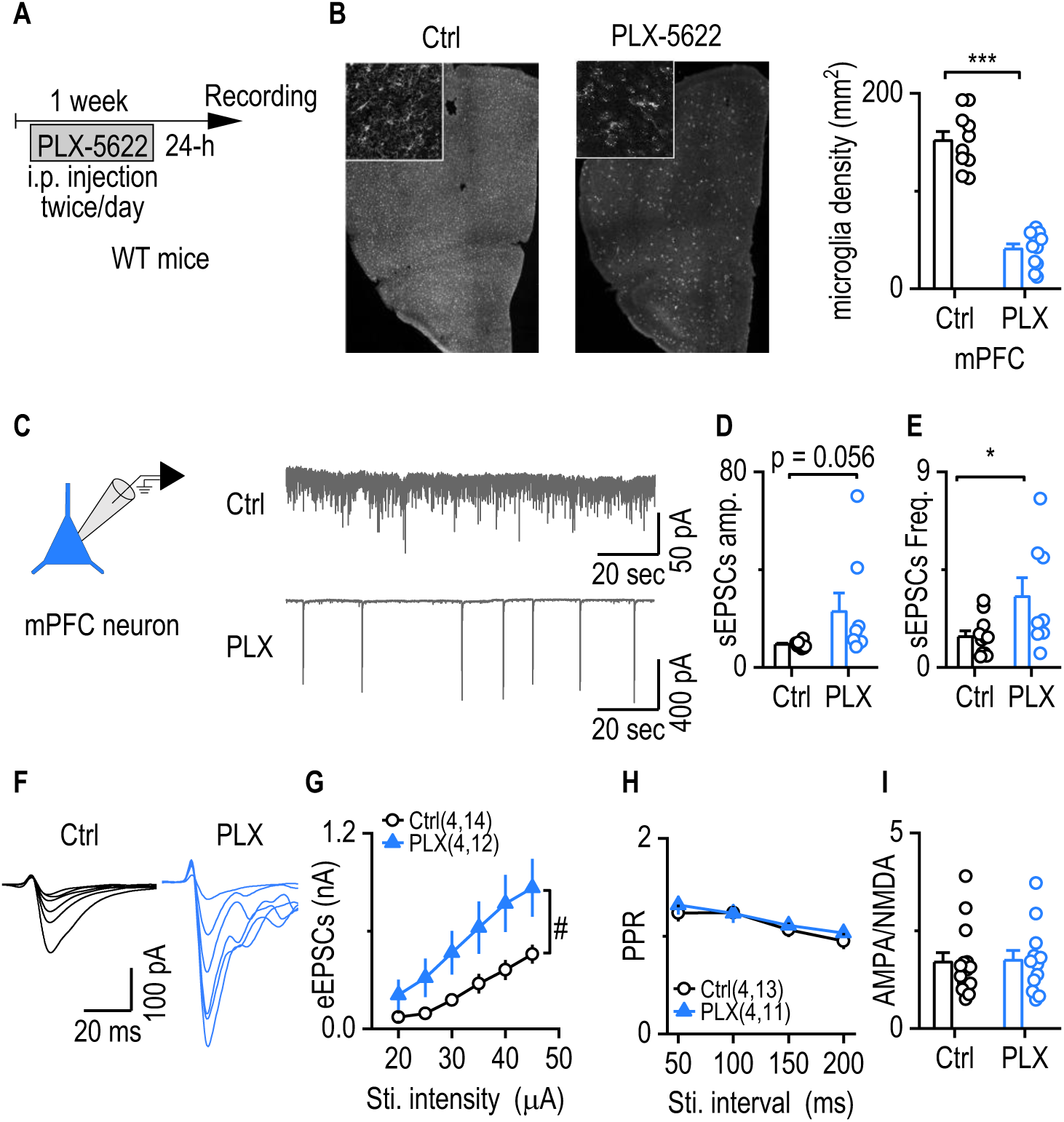
Glutamatergic transmission is enhanced under microglia-deficient conditions in WT mice. **A**, Experimental timeline of PLX5622 administration. Wild-type (WT) mice were injected intraperitoneally with PLX5622 (50 mg/kg) twice daily. Electrophysiological recordings were performed on mPFC slices prepared 24 hours after the final injection. **B,** Representative images showing IBA1+ microglia in the mPFC of saline- and PLX5622-injected mice. PLX5622 treatment significantly reduced microglial expression levels compared to saline controls. Unpaired t-test, \*\**p* < 0.01. n = 3 mice from each group. **C,** Representative traces of spontaneous excitatory postsynaptic currents (sEPSCs) recorded from mPFC neurons in both groups. **D,** sEPSC amplitudes showed a trend toward being higher in PLX5622-injected mice compared to controls. Mann Whitney test, \**p* < 0.05. n = 11 neurons from 3 mice (11/3) for Ctrl and 8/3 for PLX. **E**, sEPSC frequency was significantly increased in PLX5622-injected mice. Unpaired t test with Welch’s correction, \**p* < 0.05. n = 11/3 (Ctrl) and 8/3 (PLX). **F**, Representative traces of electrically evoked EPSCs (eEPSCs) in saline- and PLX5622-injected mice. **G,** eEPSC amplitudes were significantly higher in PLX5622-injected mice than in saline-injected controls. Two-way RM ANOVA with Greenhouse–Geisser correction, followed by Sidak’s post hoc test; ^#^p < 0.05. n = 12/3 (Ctrl), 14/3 (PLX). **H**, Paired-pulse ratios (PPRs) of eEPSCs were not significantly different between groups. Two-way RM ANOVA with Greenhouse–Geisser correction, followed by Sidak’s post hoc test. n = 11/3 (Ctrl), 13/3 (PLX). **I,** AMPA/NMDA ratios were also comparable between groups. Unpaired t-test, \**p* < 0.05. n = 14/3 (Ctrl), 12/3 (PLX).

PLX5622 treatment significantly reduced microglial density in the mPFC compared with saline-injected controls (Fig. 6B, t_4_ = 5.371, *p* = 0.0058). Microglial depletion led to a significant increase in the amplitude (Fig. 6C and D, *U* = 17, *p* = 0.0259) but not the frequency (Fig. 6E, t_2.015_ = 8.303, *p* = 0.0773) of spontaneous excitatory postsynaptic currents (sEPSCs) in mPFC neurons. Similarly, the amplitude of eEPSCs was significantly elevated in PLX5622-treated mice (Fig. 6F and G, *F*_(1,24)_ = 4.805, *p* = 0.0383), while paired-pulse ratios (Fig. 6H, *F*_(1,22)_ = 0.3959, *p* = 0.5357) and AMPA/NMDA ratio remained unchanged (Fig. 6I, *U* = 76, *p* = 0.7045).

Together, these findings support a homeostatic role for microglia in constraining glutamatergic activity in the adult brain.

## DISCUSSION

In this study, we investigated how chronic alcohol consumption differentially influences neuronal, immune, and pathological features of AD in two distinct mouse models: hAPP-KI (Aβ-driven) and PS19 (tau-driven). We found that alcohol exacerbated Aβ pathology and enhanced excitatory synaptic input to mPFC neurons in hAPP-KI mice, but reduced corticostriatal transmission and impaired striatal cholinergic signaling. In contrast, alcohol increased mPFC-to-DMS glutamatergic transmission and tau phosphorylation in PS19 mice without significantly affecting local mPFC excitability or microglial activation. Notably, microglial depletion in wild-type mice phenocopied the alcohol-induced increase in glutamatergic transmission, suggesting a critical role for microglia in maintaining excitatory balance. Together, these results reveal pathology-specific effects of alcohol on brain circuit function and disease progression and identify microglia as key regulators of alcohol-induced synaptic remodeling in AD.

A key finding of this work is the distinct modulation of mPFC-to-DMS glutamatergic transmission by chronic alcohol exposure across different AD-related models. In wild-type mice, alcohol enhances corticostriatal excitatory drive, consistent with prior reports that alcohol facilitates corticostriatal circuit activity and reward-related behavior (Wang et al., 2015; Cheng et al., 2017; Ma et al., 2017; Ma et al., 2018; Xie et al., 2023). Similarly, 5xFAD mice that modeling early and aggressive Aβ pathology exhibit elevated mPFC-to-DMS transmission during early disease stages and maintain heightened corticostriatal activity even at 12 months of age compared with aged WT mice (Huang et al., 2025), suggesting that Aβ pathology may potentiate corticostriatal hyperconnectivity. In contrast, in hAPP-KI mice, which display slower and more physiologically relevant Aβ accumulation, chronic alcohol intake significantly reduced mPFC-to-DMS transmission. This divergence may be explained by the pronounced microglial recruitment we observed in alcohol-exposed hAPP-KI mice, particularly around Aβ plaques. Because overactivated microglia can release proinflammatory cytokines, promote synaptic pruning, and induce neuronal apoptosis (Badimon et al., 2020; Lv et al., 2024). It is plausible that chronic alcohol exacerbates Aβ-driven neuroinflammation, leading to degeneration or silencing of corticostriatal projection neurons.

In contrast, our AAV-induced tau expression experiments demonstrated that tau pathology alone is sufficient to reduce mPFC-to-DMS synaptic strength, in line with previous studies reporting tau-mediated synaptic and axonal dysfunction (Busche et al., 2019; Busche and Hyman, 2020; Harris et al., 2025). Notably, in the PS19 tauopathy model, chronic alcohol consumption restored corticostriatal excitatory transmission despite no significant changes in local mPFC excitability or microglial activation. This reversal may reflect alcohol’s direct modulation of tau phosphorylation or synaptic plasticity mechanisms independent of immune activation. Indeed, we observed increased phospho-tau levels in both the mPFC and DMS following alcohol exposure, suggesting that alcohol may alter tau pathology in a manner that paradoxically enhances long-range synaptic output. These results emphasize the importance of pathological context in shaping alcohol’s effects on neural circuits and reveal distinct circuit vulnerabilities in Aβ-versus tau-driven AD models.

The observed reduction in striatal ACh release dynamics in alcohol-exposed hAPP-KI mice may stem from disrupted excitatory input to cholinergic interneurons (CINs), which require excitatory inputs for normal function. The decrease in mPFC-to-DMS transmission in this group likely contributes to reduced glutamatergic input to the striatum, leading to diminished CIN activation and impaired cholinergic output. However, corticostriatal projections are not the sole excitatory input to CINs. Thalamostriatal projections, particularly from the parafascicular and intralaminar nuclei, provide major glutamatergic input to CINs and are known to degenerate in AD (Matamales et al., 2016; Ma et al., 2022; Huang et al., 2024). Aβ accumulation in the thalamus has been reported in both patients and animal models, and chronic alcohol consumption may exacerbate this vulnerability. Alcohol may enhance microglial activation in the thalamus, driving neuroinflammation and neuronal loss, thereby weakening thalamostriatal input to CINs. Such disruption could compound CIN dysfunction and contribute to the observed reduction in ACh signaling in the striatum.

To further dissect the interaction between Aβ and tau pathology under alcohol exposure, we examined mPFC-to-DMS transmission in 5xFAD;Tau mice. In these double-transgenic animals, alcohol-exposed mice exhibited significantly reduced corticostriatal excitatory transmission compared with alcohol-exposed PS19 mice, suggesting that Aβ pathology alters the circuit-level response to alcohol in tauopathy. One potential mechanism involves microglial activation. In our study, robust microglial activation occurred in the mPFC of alcohol-drinking hAPP-KI mice but not in PS19 mice, implicating Aβ pathology as a driver of neuroimmune responses to alcohol. We speculate that in 5xFAD;Tau mice, Aβ-driven microglial overactivation may dominate the circuit remodeling outcome, leading to synaptic loss or functional suppression of corticostriatal projections and overriding any alcohol-induced excitatory enhancement observed in tau-only models. These results point to a potential hierarchical interaction between Aβ-induced neuroinflammation and tau-mediated synaptic dysfunction in shaping the brain’s response to chronic alcohol exposure.

Overall, our study highlights how the interplay between alcohol exposure and underlying AD pathology shapes neural circuit function and neuroimmune responses in a context-dependent manner. Rather than exerting uniform effects, alcohol engages distinct mechanisms in Aβ-versus tau-related models, with microglial activation emerging as a key factor modulating circuit vulnerability in the presence of Aβ. These findings underscore the importance of considering pathological background when evaluating environmental risk factors such as alcohol and suggest that targeting neuroimmune interactions may represent a strategy to mitigate alcohol-related exacerbation of AD progression.

## MATERIALS AND METHODS

### Animals

hAPP-KI (stock no. 034711), 5xFAD (stock no. 034848), and Tau P301S (PS19; stock no. 008169) mice were obtained from the Jackson Laboratory and subsequently backcrossed to the C57BL/6J strain. Breeding pairs were established in-house to generate the required experimental genotypes. Animals were maintained in groups under controlled conditions (ambient temperature: 23 °C; 12 h light/dark schedule with lights on at 23:00). Food and water were available ad libitum. Both sexes were included in all experiments. All procedures were approved by the Institutional Animal Care and Use Committee and carried out in accordance with the National Research Council’s Guide for the Care and Use of Laboratory Animals.

### Confocal imaging and cell counting

Following transcardial perfusion with 4% paraformaldehyde (PFA) in phosphate-buffered saline (PBS), brains were collected, post-fixed overnight in 4% PFA/PBS, and cryoprotected in 30% sucrose. Serial coronal sections (50 µm) were prepared using a cryostat. Confocal images were captured with an Olympus Fluoview 3000 laser-scanning microscope. Fluorescent image stacks were reconstructed in three dimensions, and manual cell counts were performed using Bitplane Imaris software (version 8.3.1; Bitplane, Zurich, Switzerland), as described previously (Wei et al., 2018). The Imaris Spot module was used for identifying neurons or microglia and for assessing colocalization. Cell density was calculated by quantifying labeled neurons or microglia within a defined circular region of interest (ROI) in the target brain area and dividing the count by the ROI area. All quantifications were conducted with experimenters blinded to group identity. Anatomical boundaries were determined using the Paxinos and Franklin mouse brain atlas (Franklin and Paxinos, 2007).

### Locomotor activity test

The open-field test was conducted as previously described (Wu et al., 2020). A transparent, open-field activity chamber (Med Associates, 43 cm x 43 cm x 21 cm height) was equipped with an infrared beam detector connected to a computer. Animals were moved to the locomotor activity test room 30 minutes before testing. The distance traveled and velocity were analyzed using Activity Monitor software (MED Associates, St Albans, VT).

### Stereotaxic virus infusion

Stereotaxic viral infusions were performed as previously described (Wang et al., 2015; Huang et al., 2017; Ma et al., 2017; Roltsch Hellard et al., 2019). Stereotaxic viral delivery was carried out under aseptic conditions. After exposing the skull, the bregma and lambda landmarks were identified, and their spatial coordinates were recorded using a three-axis micromanipulator. Small craniotomies were drilled at stereotaxic coordinates determined from the Paxinos and Franklin mouse brain atlas. Viral solutions were infused at a rate of 0.1 µL/min. For striatal injections, 0.5 µL of AAV2/9-hSyn-gACh4m (BrainVTA, PT-7021) was delivered bilaterally (AP: +0.26 mm, ML: ± 2.00 mm, DV: −3.50 mm). For mPFC-to-DMS transmission studies, 0.5 µL of rAAV8/svn-ChR90-GFP (Svn-Chronos-GFP; UNC, AV5842B) was infused bilaterally into the mPFC. In a separate set of experiments, WT mice received bilateral mPFC co-infusions of 0.4 µL rAAV9/syn-Flex-ChrimsonR-tdT (UNC, AV6556B) with either 0.4 µL rAAV2/8-CMV-Tau(P301L)-eGFP-WPREs (BrainVTA, PT-6560) or AAV-EF1a-eGFP (Addgene, 105547). To minimize viral backflow, microinjectors remained in place for 10 min after infusion before withdrawal. The incision was sutured, and mice were allowed to recover for at least one week before behavioral or electrophysiological testing.

### PLX 5622 Preparation

The PLX5622 preparation was previously described(Gangal et al., 2025). A 5 mg/mL (12.65 mM) working solution of PLX 5622 (CHEMGOOD, C-1521) was prepared by first dissolving the compound in dimethyl sulfoxide (DMSO; Sigma, D8418) to generate a concentrated DMSO–PLX 5622 stock. The solvent for dilution consisted of 20% ethoxylated hydrogenated castor oil (MedChemExpress, HY-126403) in saline. The stock solution was then diluted into this solvent to achieve a final mixture containing 5% DMSO– PLX 5622 and 95% of the 20% ethoxylated hydrogenated castor oil in saline. The resulting solution was ultrasonicated until fully transparent.

### Electrophysiological recordings of brain slices

#### Brain slice preparation

Immediately after brain extraction, 250-µm coronal sections containing either the striatum or basal forebrain were prepared in ice-cold cutting solution. The cutting solution contained (in mM): 40 NaCl, 143.5 sucrose, 4 KCl, 1.25 NaH_2_PO_4_, 26 NaHCO_3_, 0.5 CaCl_2_, 7 MgCl_2_, 10 glucose, 1 sodium ascorbate, and 3 sodium pyruvate (pH 7.35; osmolarity 305–310 mOsm), and was continuously bubbled with 95% O_2_ + 5% CO_2_. Slices were then transferred to a 32 °C chamber containing a 1:1 mixture of cutting and external solutions for 45 min. The external solution (pH 7.35; osmolarity 305–310 mOsm) consisted of (in mM): 125 NaCl, 4.5 KCl, 2 CaCl_2_, 1 MgCl_2_, 1.25 NaH_2_PO_4_, 25 NaHCO_3_, 15 sucrose, and 15 glucose, and was also saturated with 95% O_2_ + 5% CO_2_. After incubation, slices were maintained in external solution at room temperature until use.

#### Electrophysiological recordings

Brain slices were transferred to a recording chamber mounted on the fixed stage of an upright microscope (Olympus) and continuously superfused with oxygenated external solution at 32 °C (flow rate: 2 mL/min). Neurons were visualized through a 40× water-immersion objective coupled to an infrared-sensitive CCD camera. Recordings were performed using a Multiclamp 700B amplifier, Clampex 10.6 software, and a Digidata 1550A digitizer (Molecular Devices, Sunnyvale, CA). Patch pipettes (3–6 MΩ) were pulled from borosilicate glass capillaries (World Precision Instruments, Sarasota, FL) using a P-97 micropipette puller (Sutter Instrument Co.).

For electrically evoked EPSC recordings in cortical neurons or medium spiny neurons (MSNs), a bipolar glass stimulating electrode was placed in the same compartment, delivering brief current pulses (40 mA) at 0.05 Hz. Patch electrodes were filled with a cesium-based internal solution containing (in mM): 119 CsMeSO_4_, 8 TEA.Cl, 15 HEPES, 0.6 ethylene glycol tetraacetic acid (EGTA), 0.3 Na_3_GTP, 4MgATP, 5 QX-314.Cl and 7 phosphocreatine (pH 7.3, adjusted with NaOH). For optically evoked EPSCs (oEPSCs), 2 ms light pulses were delivered through the objective lens at 470 nm (Chronos) or 590 nm (Chrimson).

To assess AMPAR/NMDAR ratios, AMPAR-mediated EPSCs were measured at a holding potential of −70 mV. NMDAR-mediated EPSCs were recorded at +40 mV, measured 30 ms after the AMPAR peak to minimize AMPAR contributions. The ratio was calculated as AMPAR peak amplitude divided by NMDAR amplitude.

Whole-cell current-clamp recordings were used to measure evoked action potential firing, with electrodes filled with a potassium-based internal solution containing (in mM): 123 potassium gluconate, 10 HEPES, 0.2 EGTA, 8 NaCl, 2 MgATP, and 0.3 NaGTP (pH 7.25, 280 mOsm). Unlike whole-cell voltage clamp, cell-attached patch recordings were used to monitor spontaneous firing without holding potential when seal resistance exceeded 1 GΩ. Evoked firing was induced by injecting current steps from 0 to 500 pA in 1 s increments.

Paired-pulse ratios (PPRs) were determined by delivering two optical or electrical stimuli at fixed intervals (50–100 ms, depending on neuron type and protocol). The amplitudes of the first (P1) and second (P2) postsynaptic responses were measured, and PPR was calculated as P2/P1. At least three trials per condition were averaged for analysis.

Data were analyzed in Clampfit (pClamp 10.7, Molecular Devices), except for sEPSC analysis, which was performed using Mini Analysis software.

#### Immunohistochemistry

Aβ immunostaining was conducted as previously described (Jagirdar et al., 2021). Antigen retrieval was achieved using 90% formic acid (Sigma, SHBQ5806). Sections were blocked with 10% bovine serum albumin (BSA) (Sigma-Aldrich, A8022) to reduce non-specific binding. Free-floating cortical sections (50 μm) were incubated overnight at 4 °C with rabbit anti-Aβ (Invitrogen, 71-5800; 1:300 in 5% BSA with 0.5% PBS-Triton™ X-100 [Fisher BioReagents, BP151-500]; PBS-TX). The following day, sections were incubated for 2 h at room temperature with Alexa Fluor 647–conjugated donkey anti-rabbit IgG (H+L) (Invitrogen, A-31573; 1:500 in 5% BSA with 0.5% PBS-TX), followed by three washes in 0.5% PBS-TX.

For microglia immunostaining, sections were blocked with 10% BSA and incubated overnight at 4 °C with goat anti-IBA1 (FUJIFILM, 011-27991; 1:500 in 5% BSA with 0.5% PBS-TX). Sections were then incubated for 2 h at room temperature with Alexa Fluor Plus 488– or 647–conjugated donkey anti-goat IgG (H+L) (Invitrogen, A-32814 [488], A-32849 [647]; 1:500 in 5% BSA with 0.5% PBS-TX) and washed three times in 0.5% PBS-TX.

For activated microglia, sections were blocked with 10% BSA and incubated overnight at 4 °C with rat anti-CD68 (BIO-RAD, MCA1957; 1:500 in 5% BSA with 0.5% PBS-TX). The following day, sections were incubated for 2 h at room temperature with Alexa Fluor Plus 488–conjugated donkey anti-rat IgG (H+L) (Invitrogen, A-21208; 1:500 in 2% BSA with 0.5% PBS-TX) and washed three times in 0.5% PBS-TX.

For phosphorylated Tau (p-Tau) immunostaining, sections were blocked with 10% BSA and incubated overnight at 4 °C with mouse anti-AT8, Biotin (Invitrogen, MN1020; 1:500 in 5% BSA with 0.5% PBS-TX). The following day, sections were incubated for 2 h at room temperature with Streptavidin, Alexa Fluor™ 647 Conjugate (Invitrogen, S21374; 1:500 in 2% BSA with 0.5% PBS-TX) and washed three times in 0.5% PBS-TX.

Quantification of Aβ plaque and activated microglia coverage (% area) was performed using ImageJ (NIH).

#### Ex vivo live-tissue confocal imaging of ACh release

Acute brain slices were placed in a custom-designed recording chamber and imaged using an Olympus FluoView FV3000 confocal microscope while being continuously perfused with ACSF equilibrated with 95% O_2_/5% CO_2_. Imaging was performed with either a 10× (NA 0.3) or 40× (NA 0.8) water-immersion objective and excitation from 488 nm and 561 nm lasers. Images were acquired at a rate of 2–3 frames/s. Acquisition parameters, including laser power, high voltage (HV), gain, offset, and aperture size, were kept constant for all experiments to allow reliable comparisons. Electrical stimulation was delivered via a glass pipette containing a tungsten filament. Fluorescence responses were quantified as ΔF/F, z-score, and half-width using MATLAB-based analysis. Custom MATLAB scripts used for these analyses are available upon request.

#### Statistical analysis

Statistical comparisons were performed using two-tailed t-tests or two-way repeated-measures (RM) ANOVA followed by Sidak’s post hoc test. Data were first examined for normality and homogeneity of variance. If these assumptions were not met, alternative tests, including the Mann–Whitney U test, Wilcoxon signed-rank test, unpaired t-test with Welch’s correction, mixed-effects models, or generalized linear mixed models (GLMM), were applied as appropriate. For two-way RM ANOVA, Greenhouse–Geisser correction was used when the assumption of sphericity was violated. A significance threshold of p < 0.05 was adopted for all analyses. Statistical procedures were carried out using SigmaPlot, GraphPad Prism, or SPSS, and results are reported as mean ± s.e.m.

## Acknowledgments

This research was supported by TRSA John P. McGovern Fellowship 2024 (Y.H) and by NIH grants U01AA025932 (J.W.), R01AA027768 (J.W.), and R01AA030293 (J.W.). We thank Dr. Jianrong Li for her technical assistance.

## Author contributions

J.W. and Y.H. conceived the project and designed the experiments. Y.H. performed the behavioral experiments and analyzed the corresponding data. Y. H., X.X., H.G., and Z.H. performed the electrophysiological experiments and analyzed the corresponding data. Y.H. and X.W. performed the histology experiments. R.C. performed the live imaging confocal experiments and analyzed the corresponding data. X.W. performed animal breeding for all experiments. J.W. and Y.H. wrote the manuscript.

## Supplementary Figures

**Supplementary Figure 1.**
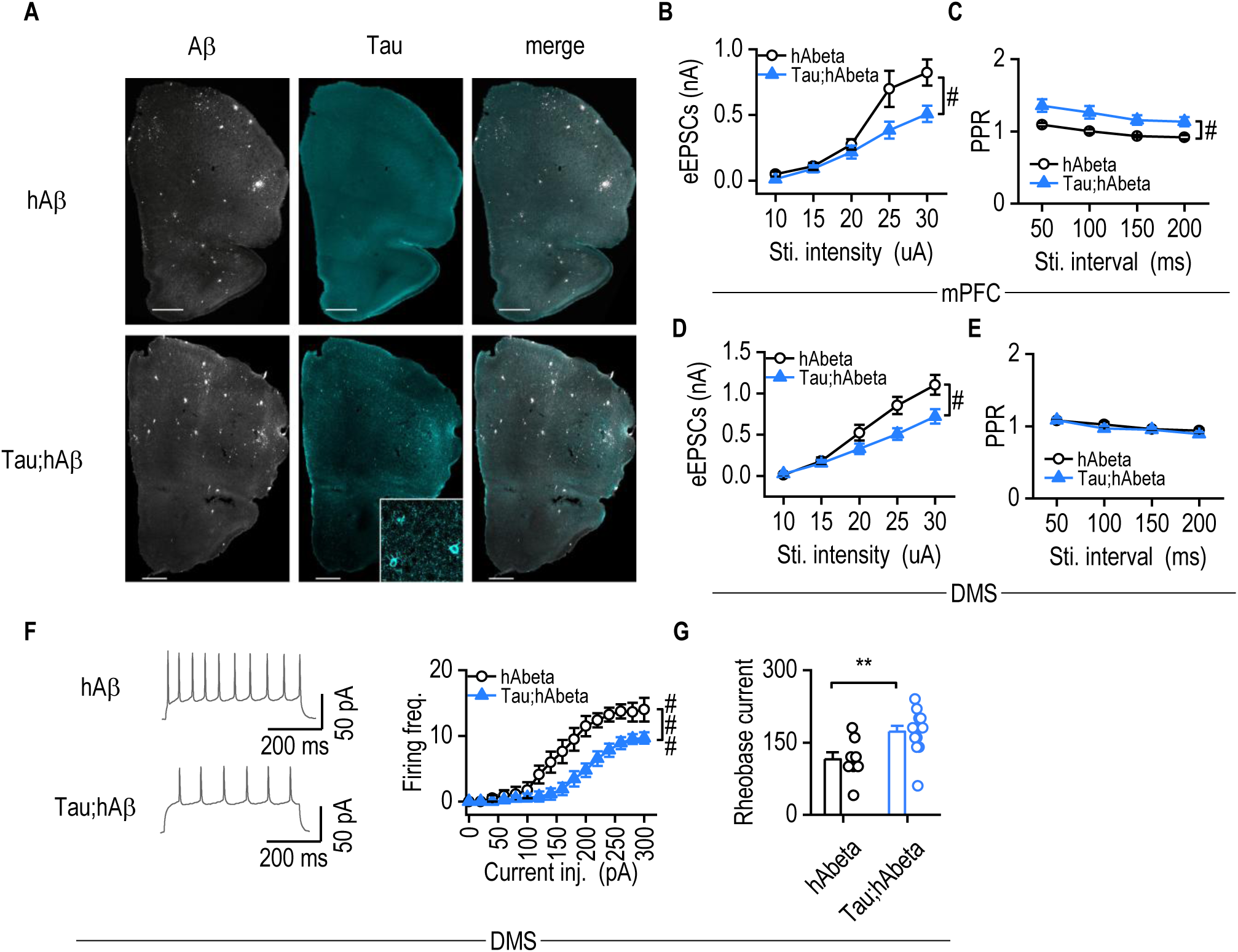
Tau expression in the hAPP-KI background reduces synaptic transmission and neuronal excitability. **A,** Representative immunostaining images showing Aβ and phosphorylated Tau (AT8) in hAPP-KI and Tau;hAPP-KI mice. **B,** Cortical neurons from Tau;hAPP-KI mice exhibited significantly lower eEPSC amplitudes compared to those from hAPP-KI mice. Two-way RM ANOVA with Greenhouse–Geisser correction, followed by Sidak’s post hoc test; ^#^*p* < 0.05. n = 10/3 for both groups. **C,** PPRs were significantly higher in cortical neurons from Tau;hAPP-KI mice relative to hAPP-KI mice. Two-way RM ANOVA with Greenhouse–Geisser correction, followed by Sidak’s post hoc test; ^#^*p* < 0.05. n = 15/3 (Tau;hAPP-KI), 10/3 (hAPP-KI). **D**, eEPSC amplitudes were also significantly lower in Tau;hAPP-KI mice than in hAPP-KI mice in DMS MSNs. Two-way RM ANOVA with Greenhouse–Geisser correction, followed by Sidak’s post hoc test; ^#^p < 0.05. n = 15/3 (Tau;hAPP-KI), 13/3 (hAPP-KI). **E,** PPRs in DMS neurons did not differ significantly between Tau;hAPP-KI and hAPP-KI mice. Two-way RM ANOVA with Greenhouse–Geisser correction, followed by Sidak’s post hoc test; ^#^p < 0.05. n = 15/3 (Tau;hAPP-KI), 14/3 (hAPP-KI). **F,** Representative traces of evoked action potentials in DMS medium spiny neurons (MSNs). Tau;hAPP-KI mice showed a lower evoked firing frequency compared to hAPP-KI mice. Two-way RM ANOVA with Greenhouse–Geisser correction, followed by Sidak’s post hoc test; ^#^*p* < 0.05. n = 13/3 (Tau;hAPP-KI), 8/3 (hAPP-KI). **G,** Rheobase current was significantly reduced in Tau;hAPP-KI mice compared to hAPP-KI mice. Unpaired t-test, \*\**p* < 0.01. n = 13/3 (Tau;hAPP-KI), 8/3 (hAPP-KI).

**Supplementary Figure 2.**
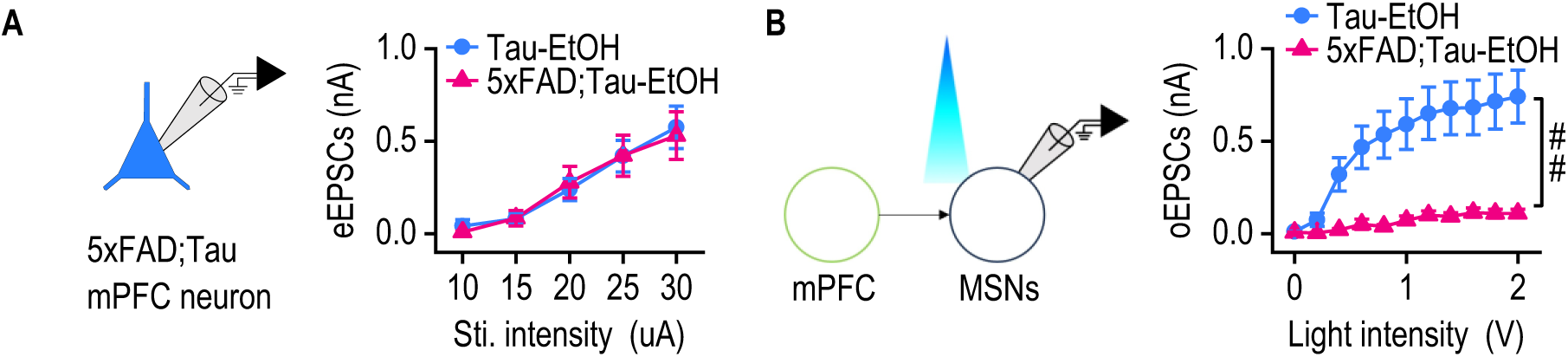
Chronic alcohol exposure reduces mPFC–DMS transmission in 5xFAD;Tau mice. **A**, In mPFC neurons, 5xFAD;Tau mice with chronic alcohol exposure exhibited eEPSC amplitudes comparable to those observed in Tau alcohol-exposed mice. Two-way RM ANOVA with Greenhouse–Geisser correction, followed by Sidak’s post hoc test. n = 10/2 (5xFAD;Tau), 12/3 (Tau). **B,** In the dorsomedial striatum, 5xFAD;Tau mice exhibited significantly lower oEPSC amplitudes in MSNs compared with Tau alcohol-exposed mice. Two-way RM ANOVA with Greenhouse– Geisser correction, followed by Sidak’s post hoc test; ^#^*p* < 0.05. n = 7/2 (5xFAD;Tau), 11/3 (Tau).

